## Supplementary Information for "Insights into glycan import by a prominent gut symbiont"

### Extended Data

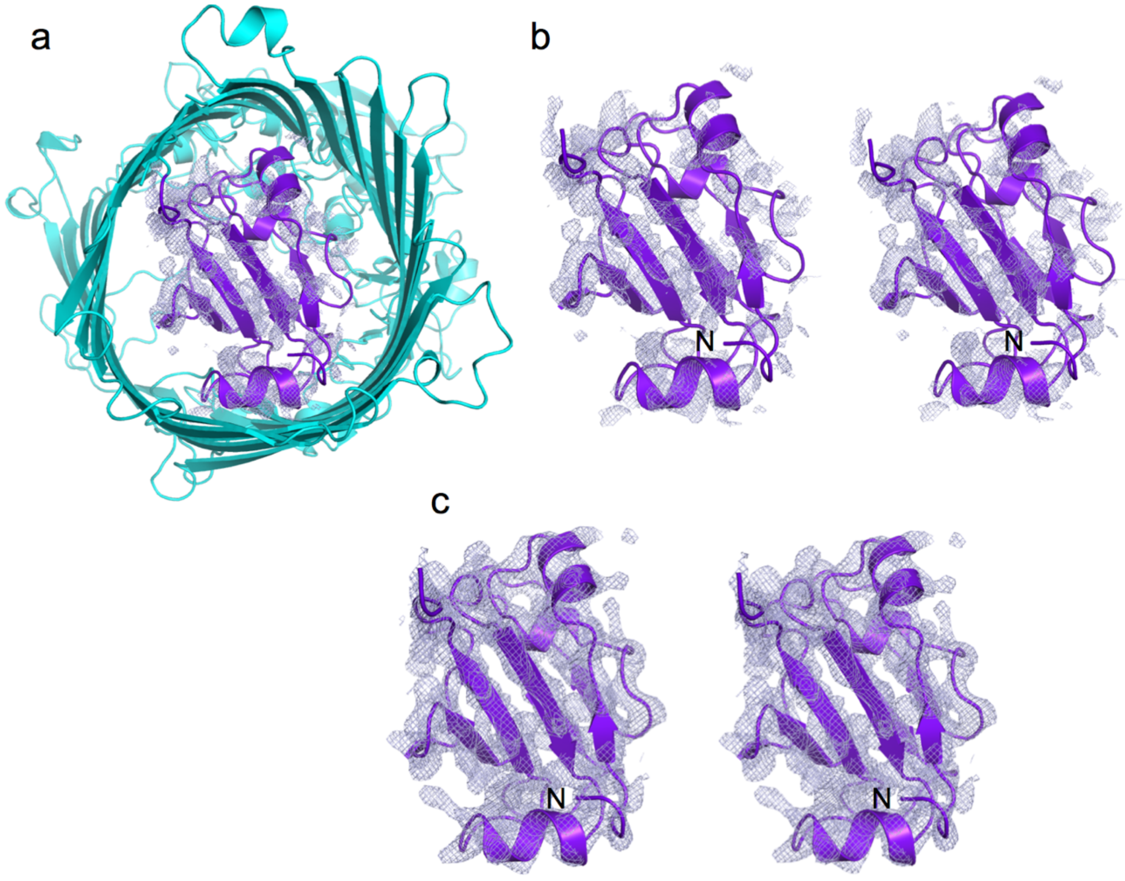

**Extended Data Figure 1** Crystal structure of apo Bt1762-63 showing poor density for the plug. **a**, Cartoon viewed from the periplasmic space, with the plug domain of the substrate-bound state superposed and coloured purple, showing weak 2Fo-Fc density for the plug ($\sigma$ = 0.8, carve = 2). **b**, Stereo cartoon with the same orientation as (**a**). The N-terminus is labelled (N). **c**, Stereo cartoon of plug density for substrate-bound Bt1762-63 ($\sigma$ = 1.5, carve = 2).

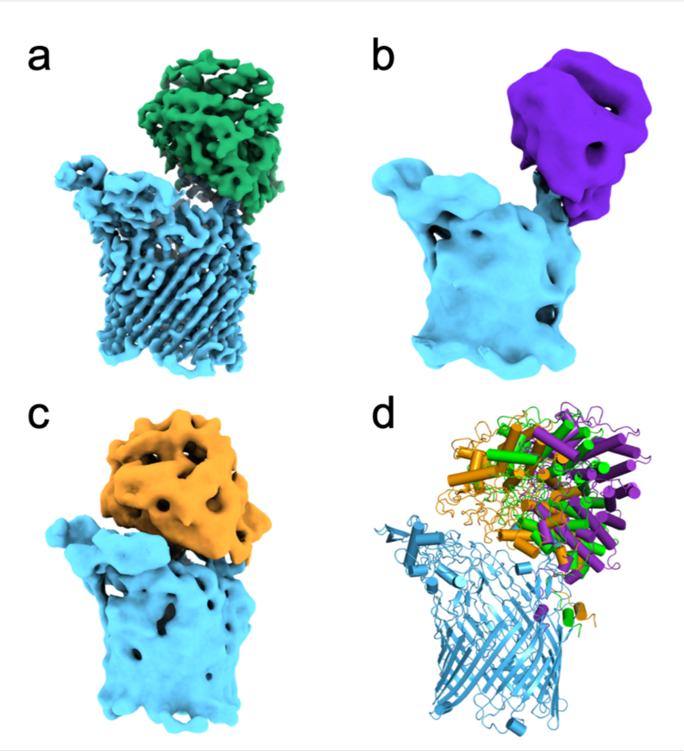

**Extended Data Figure 2** Variability in the position of Bt1762 observed in the open states of the transporter visualised by cryo-EM. **a**, Predominant open position observed in both the principle open-open and open-closed states. Less populated classes with wide open (**b**) and barely open (**c**) lid positions were also observed. **d**, Overlaid rigid-body fits of Bt1762 into the maps shown in **a-c**. EM density is filtered by local resolution and monomers are shown for clarity. The number of particles assigned to wide-open and barely-open states are 9,903 and 12,637 respectively.

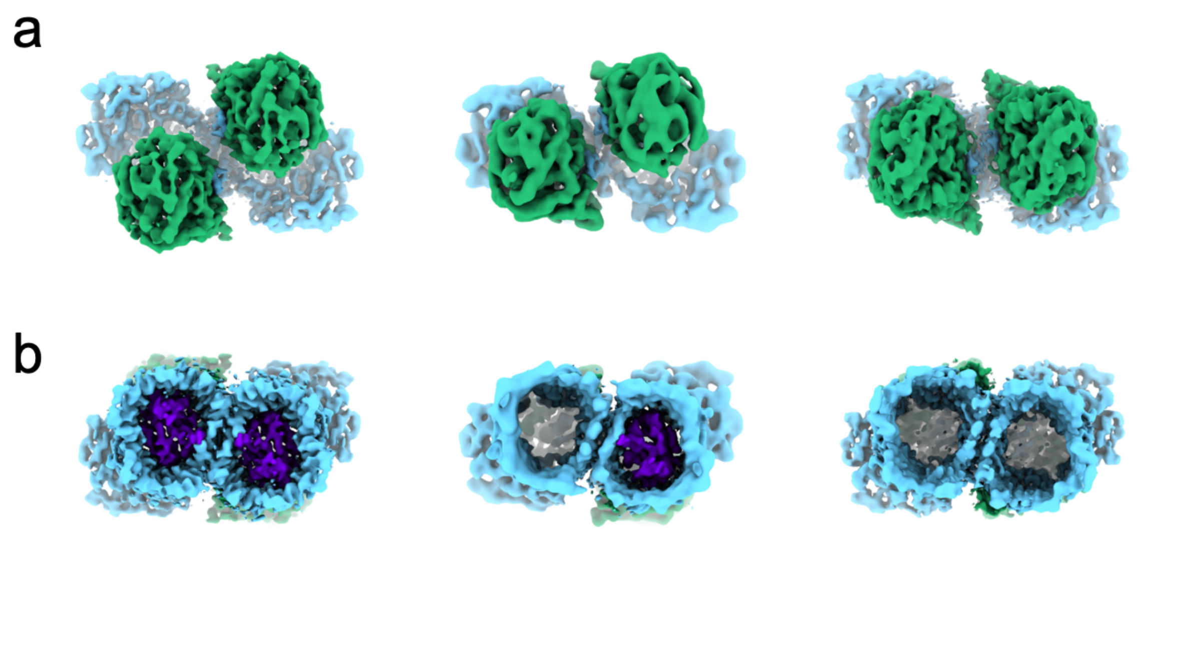

**Extended Data Figure 3** Presence or absence of the Bt1763 SusC plug domain in the three principle states from cryo-EM. Local resolution filtered EM density for Bt1762-63 in the open-open (left), closed-open (middle) and closed-closed (right) states as viewed from the extracellular (**a**) and periplasmic side (**b**) of the outer membrane. Clear density for the plug domain (purple) is invariably associated with the open state of the transporter. Where the lid is closed, no density is observed within the barrel of Bt1763.

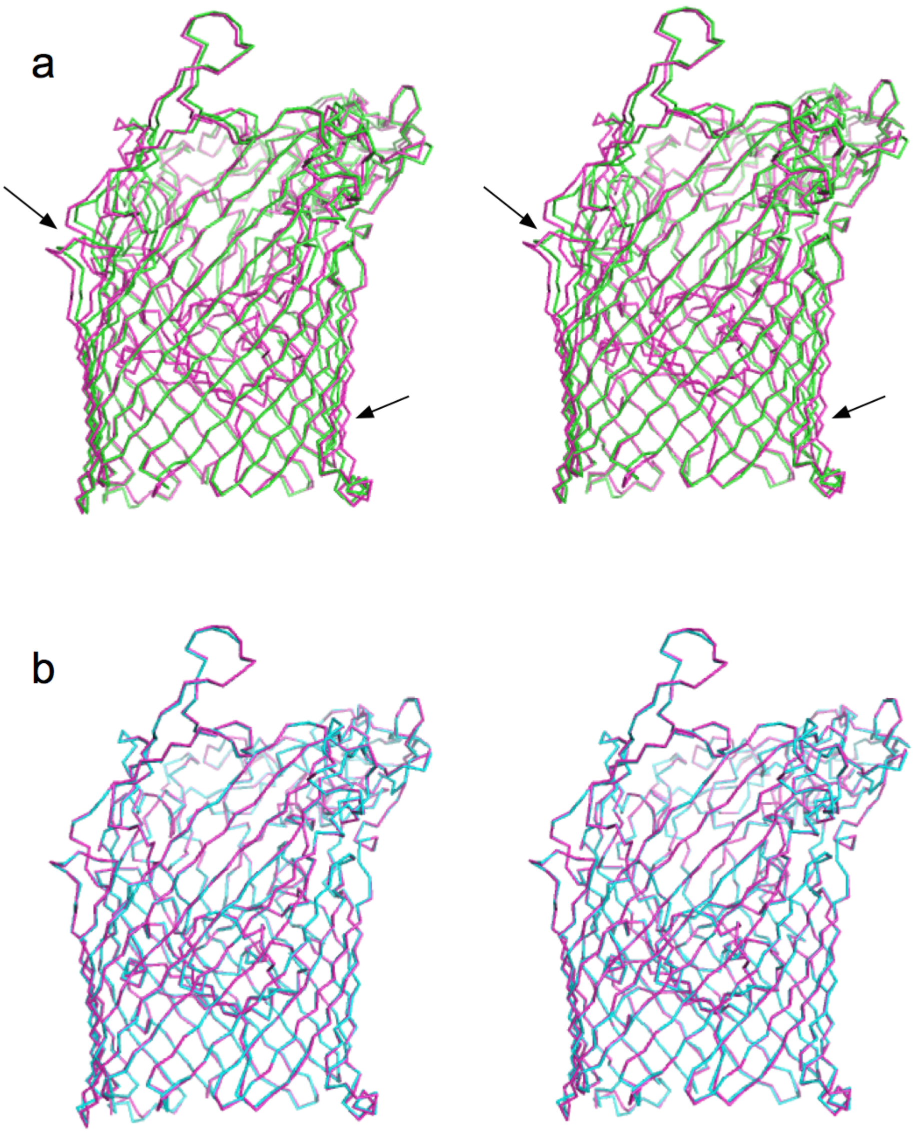

**Extended Data Figure 4** Differences in barrel shape for apo- and substrate-bound Bt1763. **a,b** Stereo ribbon superpositions for apo-Bt1763 (green; PDB 6Z8I) and substrate-bound Bt1763 (magenta; PDB 6ZAZ) (**a**) and for both substrate-bound Bt1763 structures (PDB ID 6Z9A and 6ZAZ) (**b**). Superpositions were generated in COOT via secondary structure matching for Bt1763. The arrows highlight some of the differences within the barrels. Both structures of substrate-bound Bt1763 are identical (**b**).

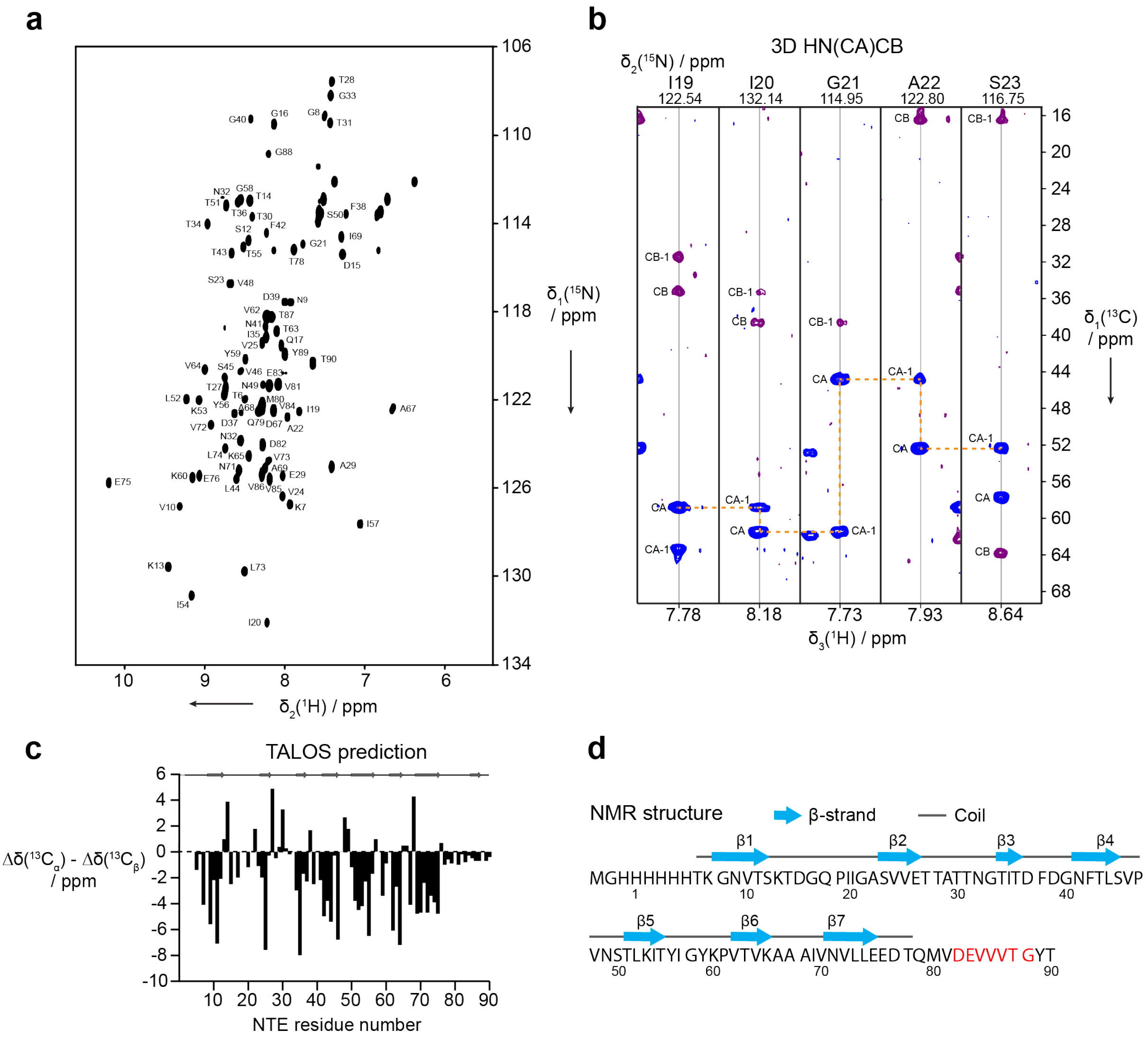

**Extended Data Figure 5** NMR backbone assignment and secondary structure of the NTE. **a**, 2D [$^{15}$N,$^{1}$H]-HSQC spectrum of NTE at 20 °C and pH 7.5. Sequence-specific resonance assignments are indicated. **b**, Example of backbone strips taken from the 3D HN(CA)CB experiments for residues I19–S23. Positive and negative intensity is shown in blue and purple, respectively. The $C_\alpha$ backbone walk is indicated by an orange dashed line. **c**, Secondary chemical shifts relative to random coil values. Consecutive stretches of negative values indicate β-sheet secondary structure. The result of a TALOS secondary structure prediction using the chemical shifts from backbone resonance assignment is plotted on top. **d**, Sequence of recombinant form of NTE domain including N-terminal $His_6$-tag. β-sheet secondary structure distributions over the NTE amino acid sequence as determined by solution NMR spectroscopy. The Ton box is shown in red.

```
# Job:
# Query: s001A
# No:  Chain   Z    rmsd lali nres  %id PDB  Description
   1:  5aq0-B 10.2  1.5   70    82   23   MOLECULE: CARBOXYPEPTIDASE D;
   2:  1uwy-A  7.9  1.7   67   393   22   MOLECULE: CARBOXYPEPTIDASE M;
   3:  5hbb-C  7.9  3.9   76   267   13   MOLECULE: CELL SURFACE PROTEIN SPAA;
   4:  4uzg-A  7.8  2.5   72   277   18   MOLECULE: SURFACE PROTEIN SPB1;
   5:  5k39-B  7.8  5.7   76   153   24   MOLECULE: CELLULOSOME ANCHORING PROTEIN COHESIN REGION;
   6:  3e8v-A  7.7  2.1   70    82   21   MOLECULE: POSSIBLE TRANSGLUTAMINASE-FAMILY PROTEIN;
   7:  6fwv-A  7.6  7.8   82   522   17   MOLECULE: COLLAGEN ADHESION PROTEIN;
   8:  4oq1-A  7.3  2.7   71   331   14   MOLECULE: CELL WALL SURFACE ANCHOR FAMILY PROTEIN;
   9:  3kpt-B  7.1  3.4   74   355   15   MOLECULE: COLLAGEN ADHESION PROTEIN;
  10:  4p0d-A  6.7  2.6   72   470   14   MOLECULE: TRYPSIN-RESISTANT SURFACE T6 PROTEIN;
  11:  1r6v-A  6.6  9.4   81   671   12   MOLECULE: SUBTILISIN-LIKE SERINE PROTEASE;
  12:  5n2j-B  6.5  2.3   73  1381   12   MOLECULE: UDP-GLUCOSE-GLYCOPROTEIN GLUCOSYLTRANSFERASE-LIKE
  13:  4fl4-I  6.3  2.3   73   309   19   MOLECULE: GLYCOSIDE HYDROLASE FAMILY 9;
  14:  4jdz-B  6.2  5.2   77   445   14   MOLECULE: SER-ASP RICH FIBRINOGEN/BONE SIALOPROTEIN-BINDING
  15:  1eo2-B  6.2  5.2   73   238   15   MOLECULE: PROTOCATECHUATE 3,4-DIOXYGENASE ALPHA CHAIN;
  16:  6fb3-A  6.2  6.9   75  1836   24   MOLECULE: TENEURIN-2;
  17:  4eiu-A  6.2  3.3   77   242   17   MOLECULE: UNCHARACTERIZED HYPOTHETICAL PROTEIN;
  18:  2y1v-A  6.1  3.5   73   604   18   MOLECULE: CELL WALL SURFACE ANCHOR FAMILY PROTEIN;
  19:  5z0z-B  6.1  2.8   73   450   16   MOLECULE: PILUS ASSEMBLY PROTEIN;
  20:  3uaf-A  6.0  4.0   70   117   20   MOLECULE: TTR-52;
  21:  3pf2-A  5.8  2.7   71   317   18   MOLECULE: CELL WALL SURFACE ANCHOR FAMILY PROTEIN;
  22:  2ww8-A  5.6  6.8   81   815   17   MOLECULE: CELL WALL SURFACE ANCHOR FAMILY PROTEIN;
  23:  5z5m-A  5.6  2.6   69   273   12   MOLECULE: PREDICTED PROTEIN;
  24:  5vxt-B  5.6  2.2   70   312   24   MOLECULE: CATECHOL 1,2-DIOXYGENASE;
  25:  4iyk-A  5.6  5.9   76   210    8   MOLECULE: UNCHARACTERIZED PROTEIN;
  26:  4z8w-A  5.5  3.0   70   125   23   MOLECULE: MAJOR POLLEN ALLERGEN PLA L 1;
  27:  6ts2-D  5.5  2.2   68  1123   13   MOLECULE: UDP-GLUCOSE-GLYCOPROTEIN GLUCOSYLTRANSFERASE-LIKE
  28:  6jch-A  5.2  2.5   70   327   17   MOLECULE: PILUS ASSEMBLY PROTEIN;
  29:  5xyr-A  5.2  2.4   62  1380    8   MOLECULE: CHEMOKINE PROTEASE C;
  30:  2x5p-A  5.2  2.4   67   104   16   MOLECULE: FIBRONECTIN BINDING PROTEIN;
```

**Extended Data Figure 6** DALI analysis of the NTE. The thirty best hits are shown.

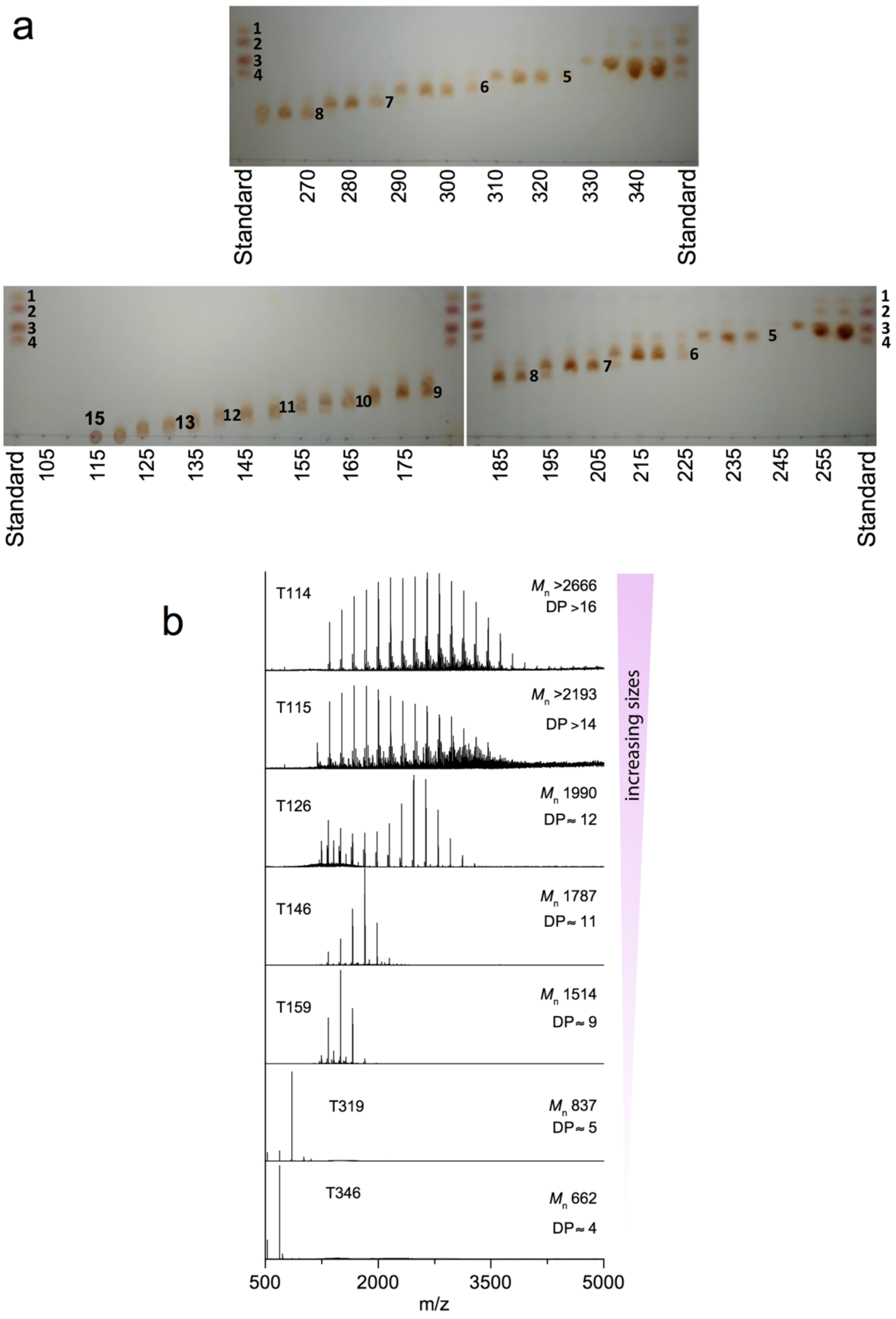

**Extended Data Figure. 7** Characterisation of β2,6 fructo-oligosaccharide (FOS) fractions. **a**, TLC analysis of digested *Erwinia* levan fractions from two different runs (top and bottom) after separation on a P2 SEC column. Approximate DP of FOS in each fraction are indicated. The standard consisted of β2,1 FOS1-4 (Fructose, sucrose, kestotriose, kestotetraose; Megazyme). **b**, Representative mass spectra of *Erwinia* FOS fractions from (**a**). The average molar mass is shown ($M_n$), based on the intensity-weighted signal intensities in the spectra.

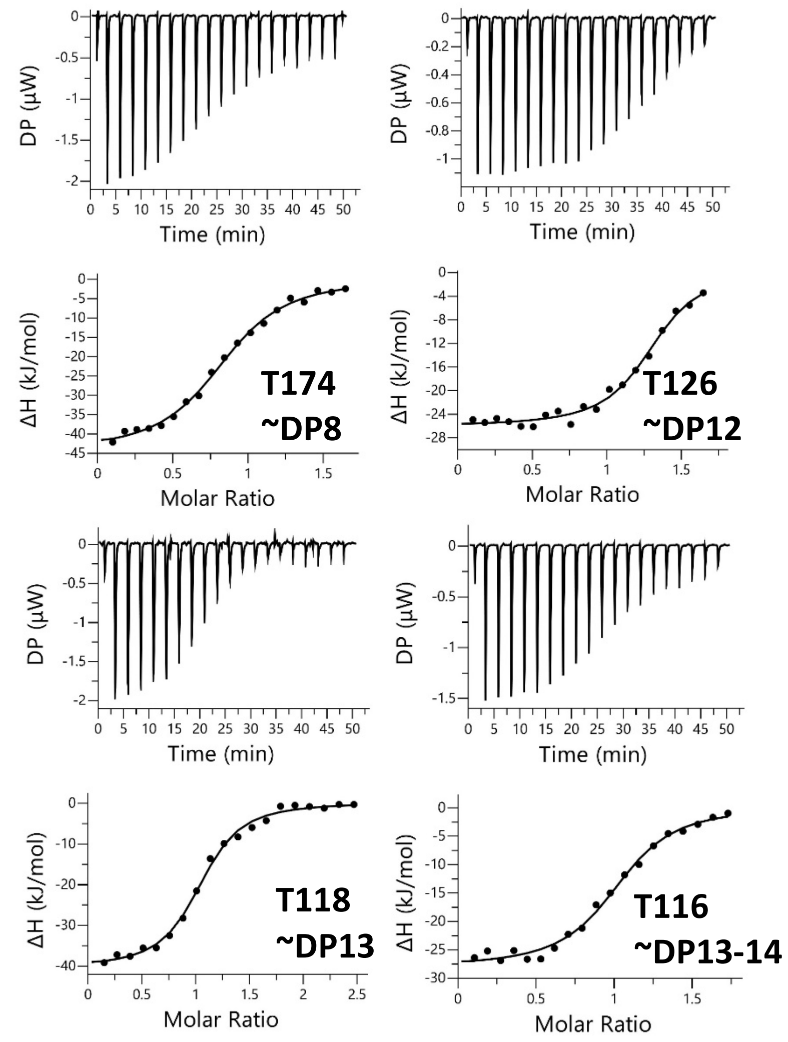

**Extended Data Figure 8** ITC traces showing FOS binding to Bt1762-63 SusCD. β2,6 FOS fractions from SEC of partially digested *Erwinia* levan were titrated into purified Bt1762-63 SusCD complex (25 μM) in 100 mM Hepes, pH 7.5 containing 0.05% LDAO. The identity of the fraction used is indicated as is the approximate DP of the main FOS species present in each fraction as determined by MS (Extended Data Fig. 7). The upper parts of each titration show the raw heats of injection and the lower parts the integrated heats, fit to a one set of sites model.

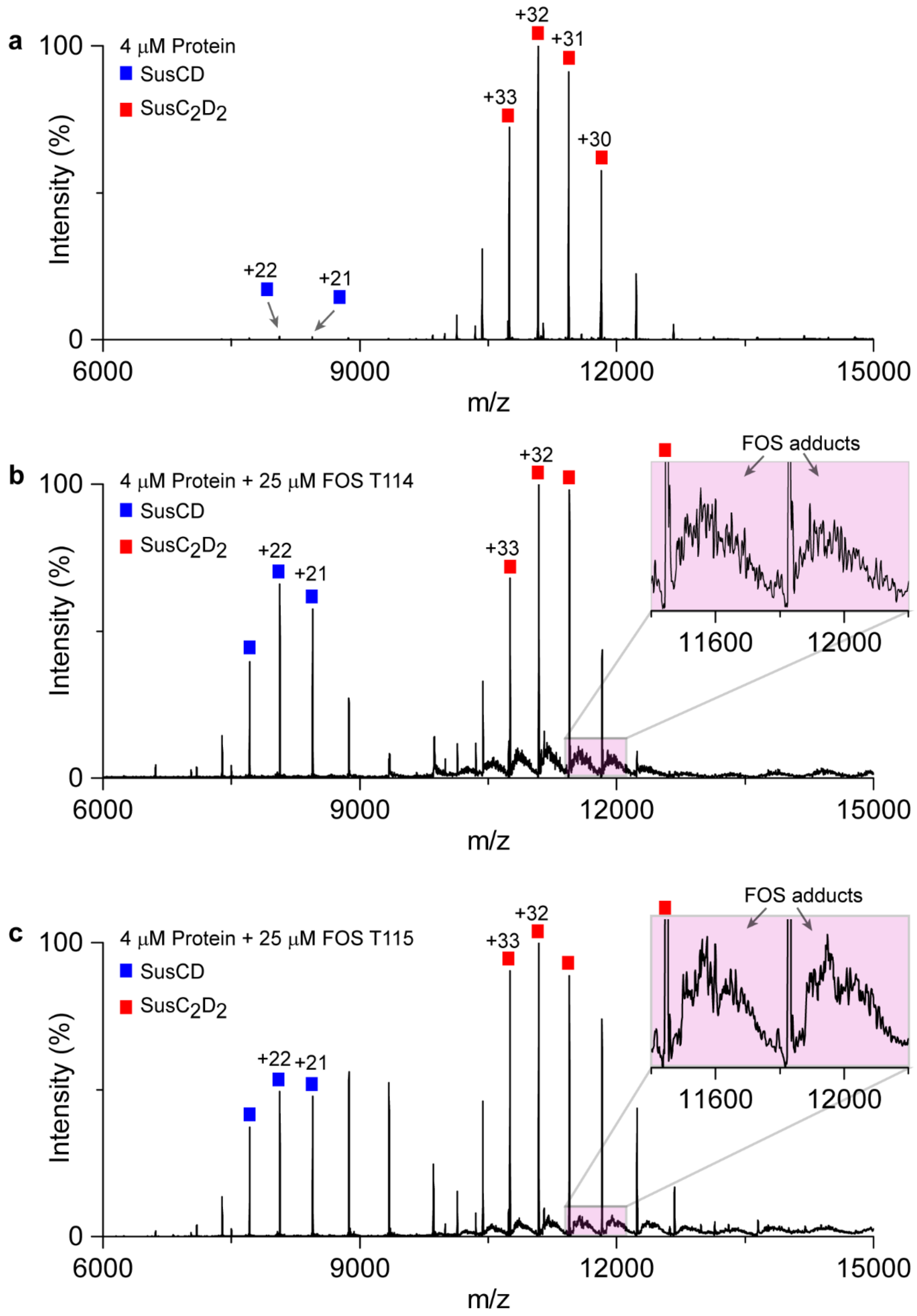

**Extended Data Figure 9** Native mass spectra of BT1762-63 without **(a)** and with long-chain FOS (**b**; T114, **c**; T115). Inserts, zoomed-in view of peaks corresponding to bound FOS molecules.

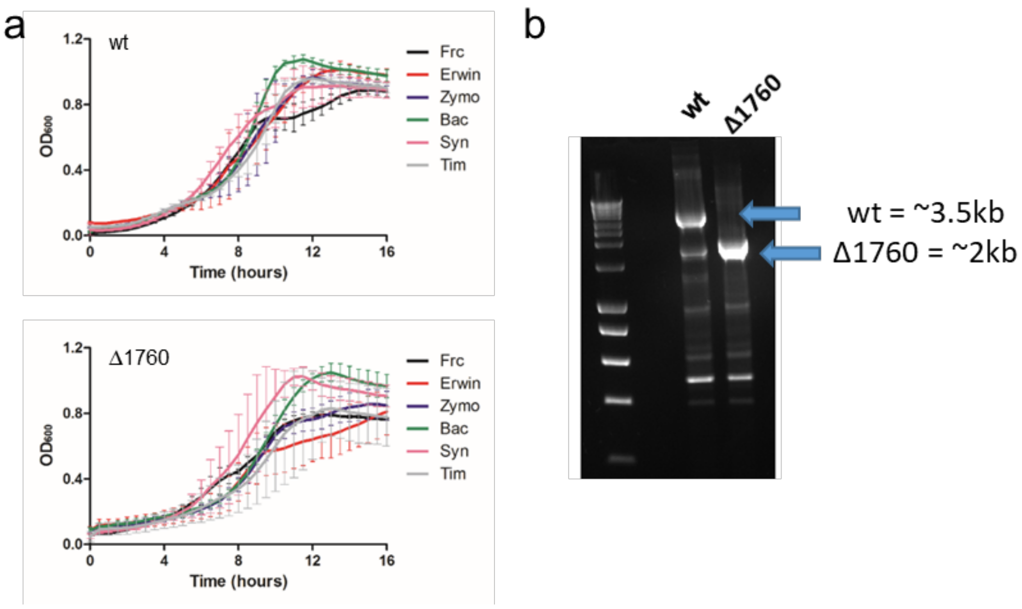

**Extended Data Figure 10** Growth of *B. theta* wild-type and Δ1760 strains on different levans. **a**, *B. theta* wild type (wt) and BT1760 deletion (Δ1760) strains were grown in minimal media with a range of levans as the sole carbon source. Levans used; Erwin = *Erwinia carotovora*, Zymo = *Zymomonas mobilis*, Bac = *Bacillus* sp., Syn = *in vitro* synthesised using levansucrase, Tim = Timothy grass. Frc = fructose. **b**, PCR analysis of genomic DNA isolated from stationary phase cells of the *Erwinia* levan cultures. Primers flanking BT1760 gene were used for amplification.

β-strand Coil

```
                      β1              β2              β3       β4
BT1763           LWGQNITVKGNVT--SKTDGQPIIGASVVETTA-----TTNGTITDFDGNFTLSVPVN-S      75
BT2264           ------------------------------------------------------------      0
PG0185 RagA      AMAQNRTVKGTVI--SSEDNEPLIGANVVVVGN-----TTIGAATDLDGNFTLSVPANAK      70
BT0268           -----------------------------------------MVIILPNFFLQKPVCRL      17
BT0272           VRAQDIPVSGVVK--DKSNSNSTLPYASVIVKNESGKTVPQLSTTTDDNGRFSTKVKSG-Y      77
BT0362           WAQDAKVLKGRI---VNAEGEPIAGAVVNVAEA-----SR-IALSDKDGFFTLKNVKPAD      82
BT3090           AFAQQITVKGHV---VDATGEPVIGASVIEGK------STNGTITDIDGNFSLNVSAN-S      76
BT3332           FVLAQVLVKGTV---KDNLGEGVPGASVQVKG------TSQGTITDLDGKFTLNIPQKNA      64
BT3680           FAQGGIDVAGIV---LDEQGQELIGVSVQIKGK-----QGVGVVTDFDGRFKITGVPAGS      69
BT3702           AFAQQITVKGIV---KDTTGEPVIGANVVVKG------TTTGTITDFDGNFQLSAKQG-D      73
BT3983           STQQQQKVTGKV---VDANNEPLIGVSVLEKG------TTNGTITDFDGNYTLVVTGSNA      171
BT4114           VSAQTITLNGNV---KDTTGEPIIGASIVEKGN-----TTNGTITDLDGNFSLKVPAN-A      73
BT4121           ANAQTRKVTGQI---VDESGQPIIGATIRLQD------ATTGTITDIDGHFSLNVPDG-K      81
BT4164           AGQVQKVISGTVTELFGKTAEPLVGVNVNLVNNQNR--SLGGGITNLNGQYNVKVPEGEK      80
BT4168           WGQGGQKLTGKI---LSATNQPVEGAIVTVLDT-------MNVTTNKEGAFQFEVKDLSK      71
BT4660           LHAQNATVKGVI---VDETDTPLIGATVQVKG------TATGSITDFDGNYTIKANKG-A      76
BT4671           AQQGGKKMTGQV---IDENKEPMIGVSILIVG------TSTGTVTDFDGNYTLNVPKDSK      80
BACOVA_02096     IQQQSVKIKGRV---TDASGEPVIGANVVQKG------TTNGTITDLNGDFTLTVSQG-A      86
BACOVA_02652     APSQNLKVSGIVT--SATDGEPLIGVSVQVKG------TSTGTITDLNGKYTLNVSTG-Q      83
BACOVA_02742     IQQQSNKVTGKV---SD-ATGPIIGASVVEKG------TANGTITDLDGNFSLNVKAG-A      85
BACOVA_03426     SAQKGITVRGTV---LDSNGETIIGASVTLKGN-----NSVGTISDIDGNFVLTVPSEKS      79
BACOVA_03428     ------------------------------------------------------------      0
BACOVA_04393     --AQTVSVTGVV---KDASGEPIIGASVVEAG------TTNGIVTDLDGNFKLNVSAK-G      73
BACOVA_04505     LWGQNITVKGNVT--SKTDGQPIIGASVVQNDS-----KSTGTITDLDGNFTLSVPTN-A      75

                   β5        β6           β7
BT1763           --TLKITYIGYKPVTVKA---AA---IVNVLL-EEDTQMVDEVVVTGYTTQR-KA-----      120
BT2264           ------------MQTQEV-AIKP---NLKVVL-RSDAQQIDEVVVTAMGIKRSEK-----      38
PG0185 RagA      --MLRVSYSGMTTKEVAI---AN---VMKIVL-DPDSKVLEQVVVLGYGTGQKLS-----      116
BT0268           --LAVVGFLGFSGAIASQSSSPANIDSVKTYM---KKNSFEKNVGSRFVTNR-SI-----      66
BT0272           --RLVFSFLAFDSLSVKVTKPSE---RMQIYL-SPTENMLDETVVVGFKRVS-KA-----      125
BT0362           --ELYVSSVGYLPATAIADF-DE---NFKIVM-DADLDEYAHTTPLPFNRKP-KK-----      129
BT3090           --ALTISFVGYKTQTVSV-NGKT---ALKVTL-QEDTEVLDEVVVVGYGTMK-KS-----      123
BT3332           --TLVISFIGYVTVEQKA-DSQK---PMVITL-KEDTKTLDEVVVVGYQEVR-RR-----      111
BT3680           --TLVFSYIGYETREIKYTATKL---KEKIAL-KEAVNEFDEVVVVGRDTQR-KV-----      117
BT3702           --IIVVSFIGYQPQELPV-AA-----QMNVIL-KDDTEILDEVVVIGYGQVK-KN-----      118
BT3983           --VLQFSYVGYQTLERAV-AGKT---AINITL-KEDAQVLDEVVVTALGIKRSEK-----      219
BT4114           --TVVISYIGMKTQEIAI-KGKS---KIDVTL-SDDAKALDEVVVIGYGTAK-RK-----      120
BT4121           --KVVISYIGYLKQVILP-K-GD---TLKVIL-QEDNQKLDEVVVVGYGSMK-QK-----      127
BT4164           DLTIVYSYIGMKTKRIKY-TGQT---LLNVTL-ESESMAVDEVVVSARRLNRNDLGISDK      135
BT4168           AGEISVWAPGYFSVKQLIRE-RS---NIVITLIPENQYKYNETMILPFRREG-EMQLE--      124
BT4660           --VITFSYIGYKTQEIKF-TGQS---PLNVKM-IPDNQTLDEVVVVGYGTMK-RS-----      123
BT4671           --ELQFSYVGYETKVITIPVNSN---VLNVQM-KSDSQVLSDVVIIGYGTQR-KS-----      128
BACOVA_02096     --VLQVSFIGYKQQEVSLKNGQA---QVTVVL-KDDAELLDEVVVVGYGTMK-KR-----      134
BACOVA_02652     --TLVFSYIGFMEQQV-V-ATKP---VINVVL-KEDTKTLDEVVVVGYGTMK-RS-----      129
BACOVA_02742     --TLVVSYVGYKSEE-VK-AGRG---PLNITL-KEDAKALDEVVVTALGIKRERK-----      132
BACOVA_03426     --VLIVSYVGMKPQEVKVSSKGM----IKVTL-EDDTKQLEEVVVVGYGQQK-KA-----      126
BACOVA_03428     -MNRKFIYIGCTVFAMSL---------LSMTGVQAQEESKDSLVNVAFGKVA-QE-----      44
BACOVA_04393     --SLKISFIGYQTQTIPV-AGKK---QFDITL-KEDAKVLDEVVVVGYGQMK-RS-----      120
BACOVA_04505     --LLAISYIGYKEVIIAA---KP---SLKIVM-EEDAKMIDEVVVTGYMAEK-KA-----      120

BT1763           DLTGAVSVVKV-DEIQK-QGENNPVKALQGRVPGMNITADGNPSGS-ATV-R------IR      170
BT2264           ALGYAATSVGG-EKIAE-SRTSDVMSSLAGKIAGVQISSTSSDPGASNSV-I------IR      89
PG0185 RagA      TVSGSVAKVSS-EKLAE-KPVANIMDALQGQVAGMQVMTTSGDPTAVASV-E------IH      167
BT0268           DSFGVSDTVDI-KMLQR-SQFLSIQQLLKGNVPGVYVQENNGEPGTIQSM-L------VR      117
BT0272           AVTASVTVIKA-EDLVN-TPVANPMELLQGRVPGLNIQMNNGTPGGLPSF-S------IR      176
BT0362           FVTESTSIVTG-EELEK-HPVTVLQNAFTSTVTGVETYEAQSEPGWSETAMY------IR      181
BT3090           DLTGAVSSVGV-KDIKD-SPVANIGQAMQGKVSGVQII-DAGKPGDNVTI-K------IR      173
BT3332           DLTGSVAKANM-ADVLT-APVASFDQALGGRIAGVNVTSGEGMPGGNMSI-V------IR      162
BT3680           SVVGAITNVDP-AGIQA--PAVSVSNMLGGRVPGIIAVTRSGEPGNNFSEFW------IR      168
BT3702           DMTGSVMAIKP-DELSK-GITTNAQDMLSGKIAGVSVISNDGTPGGGAQI-R------IR      169
BT3983           ALSYNVQQVNA-DAVTT-NKDPNFINSLSGKVAGVNINASSSGVGGVSKV-V------MR      270
BT4114           DITGSVATVNA-EALTV-VPVASATEALTGKMAGVQITTTEGSPDAEMKI-R------VR      171
BT4121           NITGSVSTISA-EELED-LPVSNLSEALQGMVNGLNVQLGSSRPGTNANEVYIRQNRTFT      185
BT4164           EMVSATQKVDMEKLIAA-APVVSIEEALQGQLGGVDIVL-GGDPGSRSAI-R------IR      186
BT4168           DYTAAT-NIAK-KDFMP--GTTKIDRALTGQVAGLQVKRSSGMPGEGSYY-N------LR      173
BT4660           DLTGSVASIAA-KDVEG-FKTSSVAGALGGQIAGVQITSTDGTPGAGFSI-N------IR      174
BT4671           DLTGSVASVGT-KDFNK-GMVSSPEELVNGKIAGVQIVNGGGSPTSVSTI-R------IR      179
BACOVA_02096     DLSGAVSQIKS-DDLMK-GNPTDLSKGLAGKIAGVQVNQSDGAPGGGISI-Q------IR      185
```

```
BACOVA_02652     DLTGSVVSVTG-DELKK-SVVTSLDQALQGRAAGVSVTQNSGAPGGGISV-S------IR     180
BACOVA_02742     ALGYGIDEVKG-EALTK-AKETNLINSMAGRVPGLVVSQTAGGPSGSTRV-I------LR     183
BACOVA_03426     SVVGAITQTSG-KTLERAGGVTSLGSALTGSLPGVITSASSGMPGAEDPQII------IR     179
BACOVA_03428     DLTHAISTVNT-SELTKKTANNN-------SLVGLESFV-----G------G------YN     79
BACOVA_04393     DLTGSVVSVND-EAIKK-SVVTSVDQVLQGRAAGVQVQANSGMPGGSSSI-R------IR     171
BACOVA_04505     SLTGSVAVVKM-KEVAD-IPTGNVMSGLQGRVAGMNVTTDGKPGGGNTDT-K------LR     171

BT1763           GIGTLN-----------N--NDPLYIIDGVPTKA--------------------GMHE-L     196
BT2264           GVSSLS-----------GT-NQPLYVVDGVPLNNSTVYSTDGL----NSGYDFGNGANAI     133
PG0185 RagA      GTGSLG-----------AS-SAPLYIVDGMQTSL--------------------DVVATM     195
BT0268           GLSSPVFSNK-----DVSS-VQPTVYLNGVPLMLENSYVYDIKQFDINPIGAAANMLAGL     171
BT0272           GVSDISVQSSGDGEFMMGL-TPPLFVVDGIPQEDVTGYDAAG-----LLSGATVSPLAMI     230
BT0362           GIRTMN-----------ASARSPLIIVDNVERD-----------------------LSFL     207
BT3090           GLGTIN-----------N--SNPLVVIDGIPT--------------------DLGLSSL     199
BT3332           GNNSLT-----------QE-NSPLFVIDGFPIED-------------------SSAASTL     191
BT3680           GMSTFG-----------AS-SSALVLIDGIEGN-----------------------INDL     193
BT3702           GGSSLN-----------AS-NDPLIVIDGLAIDNEGI-------------KGMANGLSMV     204
BT3983           GTKSIM-----------QS-SNALYVVDGVPMYSNANKVNGTE----FSSKGNTEPIADI     314
BT4114           GGGSIT-----------GD-NTPLFIVDGFPVES----------------------ISDI     197
BT4121           GISKDG-----------GN-STPLIIIDDVIQLGTNG-------------QPSMEQFNML     220
BT4164           GTSTLN-----------AS-SDPLIVIDGVPYPTEISDDFNFS----TATEEDLGALLNI     230
BT4168           GIRTLT-----------GD-NAPLIVINGVPHMPDKTPSALI-------DGFTRDIFQFY     214
BT4660           GVGTLT-----------GD-SSPLYIVDGFEVDD----------------------IDYL     200
BT4671           GGASLN-----------AS-NDPLIVLDGVPMEVGGS------------ISGGGNFLSLI     215
BACOVA_02096     GTNSFS-----------TN-SQPLYIVDGVPFDTGDTPASDT-----NNSQNKSNPLAFI     228
BACOVA_02652     GINSLN-----------G--NEPLYVIDGVAISGNTD--------------GNSSVLSSI     213
BACOVA_02742     GSTEMT-----------GN-NQPLYVVDGVPLDNTNFGSAGTN-----GGFDLGDGISSI     226
BACOVA_03426     TQSSWN-----------N--SEPLILVDGIERE-----------------------MSSV     203
BACOVA_03428     GSSLWG--------------QGPLVLVDGVPRS-----------------------ASSV     102
BACOVA_04393     GINSLN-----------AS-NEPIFVIDGVIIDGSTG-------------SGSDNALASI     206
BACOVA_04505     GITTIN-----------N--SSPLYVIDGVQTHD--------------------NVASII     198

BT1763           NGNDIESIQVLKDAASASIYGSRAANGVIIITTKQGKKG-QI-KINFDASVSASMYQSKM     254
BT2264           NPDDVANMTILKGAAATALYGSRAANGVVMITTKSGRKE-KGVGIEYNGGVQWSTVLR-L     191
PG0185 RagA      NPNDFESMSVLKDASATSIYGARAANGVVFIQTKKGKMS-ERGRITFNASYGISQILNTK     254
BT0268           DISSIESIEIIKDPLQLAKLGPLAANGAIWITTKDGYYGGE--NVSIGVSAGMAFAPSSV     229
BT0272           PLEDIANIQVLKDAAATSLYGSKGAYGVILIETKRGETA-KP-KVSYSANFVVKTPPRLR     288
BT0362           DAYPIESITILKDAAATAIYGMRGANGAVLVTTKRGETG-KT-KINFTQEVGFQTIAGIP     265
BT3090           NMADVERVDVLKDASATAIYGSRGANGVVMITSKRGAEG-AG-KVTVNANWAIQNATKVP     257
BT3332           NPSDIESLDFLKDASATAIYGARGANGVVIITTKKGKVG-RA-QLSYDGSFGVQHVTRTI     249
BT3680           DPADIESFSILKDASATAVYGTRGANGVVVVTTKRGKAG-KL-HVNFKTNATYSYSPRMP     251
BT3702           NPADIETLTVLKDASATAIYGSRASNGVIIITTKKGKNG-QAPSVSYNGSVSFSKTQKRY     263
BT3983           NPEDIESMSVLTGAAAAALYGSDAANGAIIITTKKGKEG--RVNITVNSNVEFNAPLV-M     371
BT4114           PASDIEDMTVLKDASSTAIYGSRGANGVILVTTKSGKEG-KI-SVNYNAYYSWKKMAKQL     255
BT4121           DPSEVESITVLRDAS-AAIYGSRAANGAILVKTKRGKKG-VP-VISYSGKFAVNDAVSHS     277
BT4164           SPNDIASVEVLKDASATAIWGTQGANGVLVIKTKQGTVG-KT-RFSFSSKWTMKEDPSTI     288
BT4168           HLQDIQNITILKGAE-AAMYGSMGSNGVILIETDGTASNDLETRVSYYGSYGINWNDKRM     273
BT4660           SNSDIESIEVLKDASSSAIYGARAANGVVLITTKSGKTG-RP-TITYNGSASYRKISKKL     258
BT4671           NPNDIESMTVLKDASSTAIYGSRASNGVIIITTKKGSGS-DI-KVSFQTTNSIATKTKTS     273
BACOVA_02096     NPHDIQSIDVLKDASATAIYGSRGANGVVIITTKRGEKG-NE-KVEFSANFTFSKIAKRM     286
BACOVA_02652     NPSDIVSMEILKDASATAIYGSRASNGVVLITTNQGKAS-KT-KVSYEGYYGLQQLPKKL     271
BACOVA_02742     NADDVENMSVLKGPAASALYGSRASHGVILITTKRANKD--KISVEYNGTLTFDTQLAKW     284
BACOVA_03426     DISSVENISVLKDASATAVYGVKGANGVILITTKRGKEG-KA-SVQIKANVTAKVASKLP     261
BACOVA_03428     KASEVESISMLKDAAAVVLYGSRAAKGVILITTKRGKDS--PMHIDVRANVGVNVPKAYP     160
BACOVA_04393     NPSDIVSMDVLKDASATAIYGARAANGVIMITTKRGQKG-EA-QITYDGYIGWQEMPKKL     264
BACOVA_04505     SSNDVESIQVLKDAASAAIYGAQAANGVIIITTKRAKEG-DV-KVSFDMSLTAQTFTGGI     256
```

**Extended Data Figure 11** ClustalOmega alignment of SusC orthologs from *B. theta* and *B. ovatus* with known glycan specificities*.* The RagA protein from *P. gingivalis* (Pg0285) and BT2264 SusC, a putative peptide importer[16], are shown as well. Only the first ~250 amino acids are shown for clarity, as SusCs are usually >1000 residues in length. The NTE is indicated in blue, with the secondary structure from NMR annotated. Residues corresponding to the Ton box are shown in red, and those for the plug loop in green (dark green for long plug loops and light green for short plug loops). The plug is coloured purple. X-ray crystal structures have been solved for BT1763 (no plug loop), BT2264 (long plug loop; PDB ID 5FQ8) and RagA (no plug loop; PDB ID 6SLN). The alignment shows there are three classes of SusC orthologs, with absent, intermediate and long plug loops.

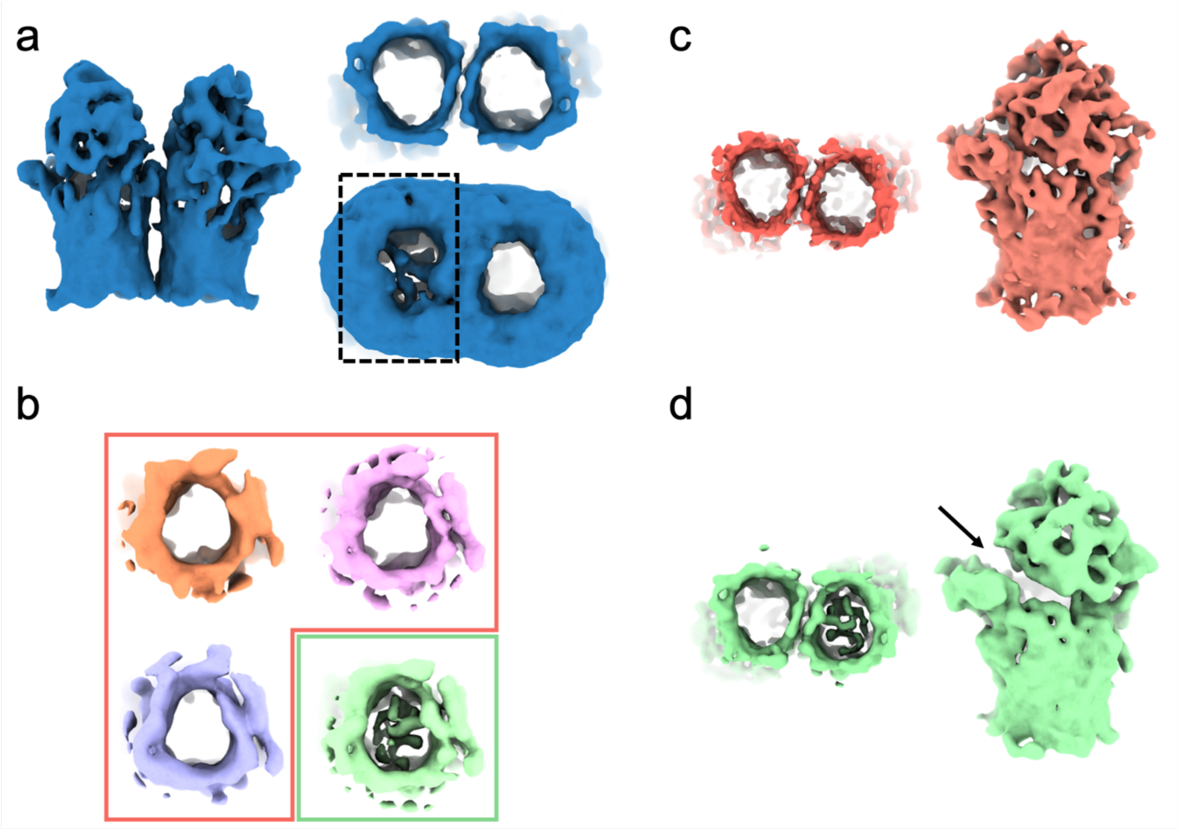

**Extended Data Figure 12** Masked classification approach and outputs. **a,** the closed-closed state observed after global 3D classification viewed in the plane of the membrane (left) and from the periplasm (right). Note that at higher contour levels, density corresponding to the plug domain can be observed in just one of the two Bt1763 barrels. Using the volume eraser tool in Chimera, a mask was generated that encompassed the plugged barrel. 'Mask create' in RELION was used to extend this mask by 5 hard and 5 soft pixels. The particle stack contributing to the reconstruction in '**a**' was subjected to 3D classification with the aforementioned mask applied. Alignment was turned off and the T value was set to 20. This classification successfully separated empty and plugged particles (**b**). Empty and plugged particles were then classified again independently in the absence of a mask. The particles contributing to the best-looking classes were refined and post-processed resulting in the reconstructions shown in **c** and **d**: A 'true' closed-closed state lacking density for the plug domain in either barrel (**c**), and a closed-barely open state where clear plug density on only one of the barrels is associated with a partially open position of the Bt1762 lid (**d**).

### Supplementary Information

**Supplementary Table 1** X-ray crystallographic data collection and refinement statistics for BT1762-63

| | apo-BT1763-63 | BT1762-63 + DP15-25 | BT1762-63 + DP6-12 |
|---|---|---|---|
| **Data collection**# | | | |
| Space group | $C222_1$ | $C222_1$ | $C222_1$ |
| Cell dimensions (Å) | | | |
| *a*, *b*, *c* (Å) | 116, 233, 171 | 120, 235,169 | 120, 238, 171 |
| $\alpha$, $\beta$, $\chi$ (°) | 90, 90, 90 | 90, 90, 90 | 90, 90, 90 |
| Resolution (Å) | 96.25-2.62 (2.66-2.62)* | 169.2-2.99 (3.04-2.99) | 171.0-2.69 (2.74-2.69) |
| $R_{pim}$ | 0.084 (1.22) | 0.13 (0.83) | 0.085 (0.61) |
| $I / \sigma I$ | 7.0 (0.8) | 5.0 (0.7) | 6.0 (1.1) |
| Completeness (%) | 99.9 (96.8) | 100 (99.3) | 100 (100) |
| Redundancy | 7.3 (7.1) | 7.2 (6.7) | 7.2 (6.9) |
| **Refinement** | | | |
| Resolution (Å) | 85.6-2.62 | 106.6-3.1 | 97.5-2.69 |
| No. reflections | 69,555 | 43,398 | 67654 |
| $R_{work}$ / $R_{free}$ (%) | 21.8/27.5 | 20.6/26.8 | 20.8/27.5 |
| No. atoms | | | |
| Protein | 10861 | 11685 | 11700 |
| ligands | - | 80 | 175 |
| Water | 80 | - | 243 |
| *B*-factors | | | |
| Protein | 66 | 70 | 49 |
| ligands | - | 77 | 65 |
| Water | 56 | - | 37 |
| R.m.s. deviations | | | |
| Bond lengths (Å) | 0.009 | 0.009 | 0.008 |
| Bond angles (°) | 1.02 | 1.19 | 1.01 |
| Clashscore | 11.5 | 13.1 | 8.2 |
| PDB ID | 6Z8I | 6Z9A | 6ZAZ |

### One crystal was used for each data collection.

* Values in parentheses are for highest-resolution shell.

**Supplementary Table 2** Cryo-EM data collection, refinement and validation statistics for Bt1762-63

| | SusCD (OO) (XXX) (XXX) | SusCD (OC) (XXX) (XXX) | SusCD (CC) (XXX) (XXX) |
|---|---|---|---|
| **Data collection and processing** | | | |
| Magnification | 130,000 x | 130,000 x | 130,000 x |
| Voltage (kV) | 300 | 300 | 300 |
| Electron exposure (e–/Å$^2$) | 63.84 | 63.84 | 63.84 |
| Defocus range (μm) | -1.5 to -3.3 | -1.5 to -3.3 | -1.5 to -3.3 |
| Pixel size (Å) | 1.07 | 1.07 | 1.07 |
| Symmetry imposed | C2 | C1 | C2 |
| Initial particle images (no.) | | 203,450 | |
| Final particle images (no.) | 32,190 | 22,205 | 17,416 |
| Map resolution (Å) FSC threshold 0.143 | 3.9 | 4.7 | 4.2 |
| Map resolution range (Å) | 3.5-5.8 | 4.3-8.3 | 3.6-7.5 |
| **Refinement** | | | |
| Initial model used (PDB code) | XXX | XXX | XXX |
| Model resolution (Å) FSC threshold (0.143/0.5) | 3.4/3.8 | 4.2/5.8 | 3.7/7.4 |
| Model resolution range (Å) | ∞ - 3.9 | ∞ - 4.7 | ∞ - 4.2 |
| Map sharpening *B* factor (Å$^2$) | -110 | -138.78 | -92.95 |
| Model composition | | | |
| Non-hydrogen atoms | 11,425 | 22,275 | 10,850 |
| Protein residues | 1,485 | 2,844 | 1,359 |
| *B* factors (Å$^2$) | | | |
| Protein (SusC/SusD) | 36.20 | 44.40 | 52.61 |
| R.m.s. deviations | | | |
| Bond lengths (Å) | 0.005 | 0.007 | 0.008 |
| Bond angles (°) | 0.867 | 0.929 | 0.988 |
| Validation | | | |
| MolProbity score | 1.93 | 2.06 | 2.08 |
| Clashscore | 4.76 | 13.46 | 17.04 |
| Poor rotamers (%) | 1.90 | 0.95 | 0 |
| Ramachandran plot | | | |
| Favored (%) | 92.30 | 93.55 | 94.91 |
| Allowed (%) | 7.70 | 6.24 | 4.65 |
| Disallowed (%) | 0 | 0.21 | 0.44 |

**Supplementary Table 3** Heteronuclear experiments and acquisition parameters used for backbone assignments and structure determination of the NTE.

| Parameter | Experiment | | | | | |
|---|---|---|---|---|---|---|
| | 2D [$^{15}$N,$^{1}$H]-HSQC | 2D [$^{13}$C,$^{1}$H]-HSQC | 3D HNCACB | 3D CBCA(CO)NH | 3D HHN NOESY | 3D HHC NOESY (ali) |
| $^{1}$H frequency (MHz) | 700 | 700 | 700 | 700 | 700 | 700 |
| $t_1$ data points | 300 | 300 | 64 | 120 | 280 | 280 |
| $t_2$ data points | 2048 | 2048 | 96 | 100 | 80 | 80 |
| $t_3$ data points | - | - | 2048 | 2048 | 2048 | 2048 |
| $t_1$ max (ms) | 52.8 | 14.2 | 13.2 | 5.6 | 90.9 | 90.9 |
| $t_2$ max (ms) | 90.9 | 90.9 | 4.5 | 17.6 | 14 | 3.78 |
| $t_3$ max (ms) | - | - | 104 | 91.7 | 12.4 | 12.4 |
| *SW* (F1) (ppm) | 40 | 60 | 34 | 60 | 16 | 16 |
| *SW* (F2) (ppm) | 16 | 16 | 60 | 40 | 40 | 60 |
| *SW* (F3) (ppm) | - | | 14 | 15.94 | 16 | 16 |
| *F1* carrier (ppm) | 117 | 45 | 120 | 45 | 4.71 | 4.71 |
| *F2* carrier (ppm) | 4.71 | 4.71 | 45 | 117 | 118 | 45 |
| *F3* carrier (ppm) | - | - | 4.74 | 4.71 | 4.71 | 4.71 |
| Interscan delay (s) | 0.8 | 0.8 | 0.8 | 0.9 | 0.8 | 0.8 |
| Number of scans | 24 | 24 | 8 | 8 | 4 | 4 |
| Mixing time (ms) | - | - | - | - | 50 | 50 |

**Supplementary Table 4** NMR structural statistics of the Bt1763 NTE.

| | NTE |
|---|---|
| **NMR distance and dihedral constraints** | |
| Distance constraints | |
| Total | 964 |
| Intra-residue (\|*i* – *j*\| = 0) | 281 |
| Inter-residue | 683 |
| Sequential (\|*i* – *j*\| = 1) | 294 |
| Medium-range 1<(\|*i* – *j*\| < 5) | 70 |
| Long-range (\|*i* – *j*\| > 5) | 319 |
| Hydrogen bonds | 0 |
| Total dihedral angle restraints | 106 |
| $\phi$ | 53 |
| $\psi$ | 53 |
| **Structure statistics** | |
| Violations (mean and s.d.) | |
| Distance constraints (Å) | 0 |
| Dihedral angle constraints (º) | 0 |
| Max. dihedral angle violation (º) | 0 |
| Max. distance constraint violation (Å) | 0 |
| Ramachandran analysis | |
| Most favored regions | 80.1% |
| Additionally allowed regions | 19.8% |
| Generously allowed regions | 0.1% |
| Disallowed regions | 0.0% |
| Average r.m.s. deviation (Å) | |
| Heavy atom | 1.03 |
| Backbone | 0.52 |

**Supplementary Table 5** Affinity of Bt1762-63 for β2,6 FOS determined by ITC.

| Ligand[a] | Kd (μM) | ΔG (kJ/mol) | ΔH (kJ/mol) | TΔS (kJ/mol) | N[b] |
|---|---|---|---|---|---|
| **DP4 (T346)** | NB[c] | - | - | - | - |
| **DP5 (T319)** | 30.5 | -25.8 | -5.3 | 20.5 | 1.4 |
| **DP6 (T305)** | 17.1 | -27.2 | -17.5 | 9.8 | 1.2 |
| **DP8 (T174)** | 1.4 | -33.4 | -44.7 | -11.3 | 0.8 |
| **DP9 (T159)** | 1.4 | -33.5 | -29.2 | 4.3 | 0.9 |
| **DP9 (T159) vs W85A[a]** | NB[c] | - | - | - | - |
| **DP11 (T146)** | 1.4 | -33.5 | -30 | 3.5 | 1.0 |
| **~DP12 (T126)** | 0.6 | -35.5 | -20.9 | 14.5 | 1.6 |
| **~DP13 (T118)** | 0.9 | -34.6 | -41 | -6.4 | 1.0 |
| **~DP13-14 (T117)** | 0.7 | -35.2 | -33.5 | 1.7 | 1.0 |
| **~DP13-14 (T116)** | 0.8 | -34.7 | -28.1 | 6.5 | 1.0 |
| **>DP14 (T115)** | 1.2 | -33.8 | -30 | 3.3 | 1.0 |
| **>DP16 (T114)** | NB[c] | - | - | - | - |
| **>DP16 (T113)** | NB[c] | - | - | - | - |

[a]DP of major species of FOS as determined by MS is shown. Fraction used (tube, T) is shown in brackets. DP4-6 are from a different FOS purification compared to the larger DP oligosaccharides. All titrations are with wild-type Bt1762-1763 complex (C-term His-tagged Bt1762), except DP9 vs W85A, which is a mutant of the Bt1762 in the complex.
[b]N is number of binding sites on the protein.
[c]NB – no binding detected.

**Supplementary Table 6** Masses of species observed in the native mass spectra shown in Fig. 7b.

| Observed | Assignment<br>Protein + adducts | Number of FOS<br>molecules bound |
|---|---|---|
| | 4uM protein alone | |
| 177405.37±0.66 | $SusC_1D_1$ | - |
| 354838.36±4.84 | $SusC_2D_2$ | - |
| | 4uM protein + 25 µM FOS 159 | |
| 177371.32±8.60 | SusCD | - |
| 178523.72±60.43 | SusCD+1152.4 | 1 |
| 178673.59±19.31 | SusCD+1301.7 | 1 |
| 178847.92±14.21 | SusCD+1476.6 | 1 |
| 179009.29±18.78 | SusCD+1638.0 | 1 |
| 356435.39±83.14 | $SusC_2D_2$+1692.75 | 1 |
| 357750.83±3.36 | $SusC_2D_2$+3008.19 | 2 |
| 357943.83±9.48 | $SusC_2D_2$+3201.19 | 2 |
| 359404.08±18.00 | $SusC_2D_2$+4661.44 | 3 |
| | 4uM protein + 50 µM FOS 159 | |
| 177380.33±14.22 | SusCD | - |
| 178544.08±1.58 | SusCD+1163.75 | 1 |
| 178697.68±7.11 | SusCD+1317.35 | 1 |
| 178861.85±6.69 | SusCD+1481.52 | 1 |
| 179023.06±14.31 | SusCD+1642.73 | 1 |
| 356515.81±27.06 | $SusC_2D_2$+1677.45 | 1 |
| 357791.44±11.15 | $SusC_2D_2$+2953.08 | 2 |
| 357980.53±6.36 | $SusC_2D_2$+3219.87 | 2 |
| 359446.38±1.34 | $SusC_2D_2$+4685.72 | 3 |
| 360665.22±171.0 | $SusC_2D_2$+5826.86 | 4 |
| | 4uM protein + 100 µM FOS 159 | |
| 177399.31±9.10 | SusCD | - |
| 178543.78±13.78 | SusCD+1163.45 | 1 |
| 178710.08±11.09 | SusCD+1329.75 | 1 |
| 178878.88±12.17 | SusCD+1498.55 | 1 |
| 179046.44±11.71 | SusCD+1666.11 | 1 |
| 180366.35±12.78 | SusCD+2986.02 | 2 |
| 180203.41±14.05 | SusCD+2823.08 | 2 |
| 180672.93±26.94 | SusCD+3292.60 | 2 |
| 357823.34±28.59 | $SusC_2D_2$+2984.98 | 2 |
| 357987.55±31.87 | $SusC_2D_2$+3149.19 | 2 |
| 359488.15±70.86 | $SusC_2D_2$+4649.79 | 3 |
| 360839.11±105.09 | $SusC_2D_2$+6001.11 | 4 |
| 361151.03±174.06 | $SusC_2D_2$+6312.67 | 4 |

**Supplementary Table 7** Primers used to construct Bt1762-63 SusCD mutants.

| Name | Sequence |
|---|---|
| TB-fwd | AAAGAAGATAACATTCGAGTCGACCTACGGGAGCATG |
| TBD-rev | CTTTACGCTGTGTAGTATAATCTACCATCTGGGTATC |
| TBD-fwd | GATACCCAGATGGTAGATTATACTACACAGCGTAAAG |
| TB-rev | GGTGGCGGCCGCTCTAGAGAGGGTTACGACGGT |
| TBM-rev | CTTTACGCTGTGTAGTATAAGCAGCAGCAGCAGCAGCATCTACCATCTGGGTATC |
| TBM-fwd | GATACCCAGATGGTAGATGCTGCTGCTGCTGCTGCTTATACTACACAGCGTAAAG |
| Hinge-fwd | AAAGAAGATAACATTCGAGTCGACACAATCTTTCCGTCAGCA |
| H2D-rev | CGAATTTATACCAGTCAGGATATCTGTTGCTTTC |
| H2D-fwd | ATCCTGACTGGTATAAATTCGGGTGCAATG |
| Hinge-rev | CACCGCGGTGGCGGCCGCTCTAGAGATCATTCGTTTTATTGA |
| H1D-rev | TTTGAAACCACCAGGTGCATAGATAGTATAACGGGC |
| H1D-fwd | TATGCACCTGGTGGTTTCAAACGCAACCAGATCGGT |
| Trp1-fwd | AAAGAAGATAACATTCGAGTCGACGAAGAGTGTAGTCGGACATAC |
| Trp2-rev | ATTGATATTCGCGTCGGTGGTATTGATACC |
| Trp3-fwd | ACCACCGACGCGAATATCAATGATATATGG |
| Trp4-rev | CACCGCGGTGGCGGCCGCTCTAGACTGAATCAGTGCTTCGGCAC |

| | |
|---|---|
| NTE1 -fwd | AAAGAAGATAACATTCGAGTCGACCTACGGGAGCATG |
| NTE2 -rev | AACTTCATCTACCATCTGAATGTTTTGCCCCCACAG |
| NTE3 -fwd | CTGTGGGGGCAAAACATTCAGATGGTAGATGAAGTT |
| NTE4 -rev | GGTGGCGGCCGCTCTAGAGAGGGTTACGACGGT |
